# Epistatic selection on a selfish *Segregation Distorter* supergene: drive, recombination, and genetic load

**DOI:** 10.1101/2021.12.22.473781

**Authors:** Beatriz Navarro-Domínguez, Ching-Ho Chang, Cara L. Brand, Christina A. Muirhead, Daven C. Presgraves, Amanda M. Larracuente

**Affiliations:** Department of Biology, University of Rochester, Rochester, New York, 14610, USA

## Abstract

Meiotic drive supergenes are complexes of alleles at linked loci that together subvert Mendelian segregation to gain preferential transmission. In males, the most common mechanism of drive involves the disruption of sperm bearing alternative alleles. While at least two loci are important for male drive— the driver and the target— linked modifiers can enhance drive, creating selection pressure to suppress recombination. In this work, we investigate the evolution and genomic consequences of an autosomal multilocus, male meiotic drive system, *Segregation Distorter* (*SD*) in the fruit fly, *Drosophila melanogaster*. In African populations, the predominant *SD* chromosome variant, *SD-Mal*, is characterized by two overlapping, paracentric inversion on chromosome arm *2R* and nearly perfect (~100%) transmission. We study the *SD-Mal* system in detail, exploring its components, chromosomal structure, and evolutionary history. Our findings reveal a recent chromosome-scale selective sweep mediated by strong epistatic selection for haplotypes carrying *Sd*, the main driving allele, and one or more factors within the double inversion. While most *SD-Mal* chromosomes are homozygous lethal, *SD-Mal* haplotypes can recombine with other, complementing haplotypes via crossing over and with wildtype chromosomes only via gene conversion. *SD-Mal* chromosomes have nevertheless accumulated lethal mutations, excess non-synonymous mutations, and excess transposable element insertions. Therefore, *SD-Mal* haplotypes evolve as a small, semi-isolated subpopulation with a history of strong selection. These results may explain the evolutionary turnover of *SD* haplotypes in different populations around the world and have implications for supergene evolution broadly.

## Introduction

Supergenes are clusters of linked loci that control complex phenotypes. Some supergenes mediate adaptive polymorphisms that are generally maintained by some form of frequency- or density-dependent natural selection, as in, *e.g*., mimicry in butterflies, self-incompatibility in plants, plumage polymorphisms in birds, and heteromorphic sex chromosomes (see Schwander *et al*. 2014; Thompson and Jiggins 2014 for review). Other supergenes are maintained by selfish social behaviors that enhance the fitness of carriers at the expense of non-carriers, as in some ant species (Keller and Ross 1998; Wang *et al*. 2013). Still other supergenes are maintained by their ability to achieve selfish, better-than-Mendelian transmission during gametogenesis, as in so-called meiotic drive complexes found in fungi, insects, and mammals (Lyon 2003; Larracuente and Presgraves 2012; Lindholm *et al*. 2016; Svedberg *et al*. 2018).

Meiotic drive complexes gain transmission advantages at the expense of other loci and their hosts. In heterozygous carriers of male drive complexes, the driver disables spermatids that bear drive-sensitive target alleles (Larracuente and Presgraves 2012; Lindholm *et al*. 2016). To spread in the population, the driver must be linked in a *cis*-arrangement to a drive-resistant (insensitive) target allele (Charlesworth and Hartl 1978). Recombination between the driver and target results in a “suicide” haplotype that distorts against itself (Sandler and Carpenter 1972; Hartl 1974). These epistatic interactions between driver and target lead to selection for modifiers of recombination that tighten linkage, such as chromosomal inversions (Schwander *et al*. 2014; Thompson and Jiggins 2014; Charlesworth 2016). Like most supergenes (Turner 1967; Charlesworth and Charlesworth 1975), meiotic drive complexes originate from two or more loci with some degree of initial linkage. Successful drivers thus tend to be located in regions of low recombination, such as non-recombining sex chromosomes (Hamilton 1967; Hurst and Pomiankowski 1991), centromeric regions, or in chromosomal inversions of autosomes (Lyon 2003; Larracuente and Presgraves 2012; Lindholm *et al*. 2016; Svedberg *et al*. 2018).

The short-term benefits of reduced recombination can entail long-term costs. Chromosomal inversions that lock supergene loci together also incidentally capture linked loci, which causes large chromosomal regions to segregate as blocks. Due to reduced recombination, the efficacy of natural selection in these regions is compromised: deleterious mutations can accumulate and beneficial ones are more readily lost (Muller 1964; Hill and Robertson 1968; Felsenstein 1974). Many meiotic drive complexes are thus homozygous lethal or sterile. The degeneration of drive haplotypes is not however inevitable, as different drive haplotypes that complement one another (Dod *et al*. 2003; Presgraves *et al*. 2009; Brand *et al*. 2015), they may be able to recombine, if only among themselves. Gene conversion from wildtype chromosomes may also ameliorate the genetic load of supergenes (Uyenoyama 2005; Wang *et al*. 2013; Tuttle *et al*. 2016; Branco *et al*. 2018; Stolle *et al*. 2019; Brelsford *et al*. 2020). Male meiotic drive complexes thus represent a class of selfish supergenes that evolve and persist via the interaction of drive, recombination, and natural selection.

Here we focus on the evolutionary genetics of *Segregation Distorter* (*SD*), a well-known autosomal meiotic drive complex in *Drosophila melanogaster* (Sandler *et al*. 1959). In heterozygous males, *SD* disables sperm bearing drive-sensitive wildtype chromosomes via a chromatin condensation defect (Hartl *et al*. 1967; Temin *et al*. 1991). *SD* has two main components: the driver, *Segregation distorter* (*Sd*), is a truncated duplication of the gene *RanGAP* located in chromosome arm *2L* (Powers and Ganetzky 1991; Merrill *et al*. 1999; Kusano *et al*. 2001); and the target of drive, *Responder* (*Rsp*), is a block of satellite DNA in the pericentromeric heterochromatin of *2R*. Previous studies of *SD* chromosomes have detected linked upward modifiers of drive, including *Enhancer of SD* (*E[SD]*) on *2L* and several others on *2R* (Sandler and Hiraizumi 1960; Miklos 1972; Ganetzky 1977; Hiraizumi *et al*. 1980; Brittnacher and Ganetzky 1984), but their molecular identities are unknown. *Sd-RanGAP* and *Rsp* straddle the centromere, a region of reduced recombination, and some *SD* chromosomes bear pericentric inversions that presumably further tighten linkage among these loci. Many *SD* chromosomes also bear paracentric inversions on *2R* (reviewed in Lyttle 1991; Larracuente and Presgraves 2012). Although recombination between paracentric inversions and the main components of *SD* is possible, their strong association implies a role for epistatic selection in the evolution of these supergenes (Larracuente and Presgraves 2012).

While *SD* is present at low population frequencies (<5%) around the world (Temin *et al*. 1991; Larracuente and Presgraves 2012), *Sd-RanGAP* appears to have originated in sub-Saharan Africa, the ancestral geographic range of *D. melanogaster* (Presgraves *et al*. 2009; Brand *et al*. 2015). The predominant *SD* variant in Africa is *SD-Mal*, which recently swept across the entire continent (Presgraves *et al*. 2009; Brand *et al*. 2015). *SD-Mal* has a pair of rare, African-endemic, overlapping paracentric inversions on *2R*: *In(2R)51B6–11;55E3–12* and *In(2R)44F3–12;54E3–10*, hereafter collectively referred to as *In(2R)Mal* (Aulard *et al*. 2002; Presgraves *et al*. 2009). *SD-Mal* chromosomes are particularly strong drivers, with ~100% transmission. Notably, recombinant chromosomes bearing the *Sd-RanGAP* duplication from this haplotype but lacking the inversions do not drive (Presgraves *et al*. 2009), suggesting that *In(2R)Mal* is essential for *SD-Mal* drive. We therefore expect strong epistatic selection to enforce the association of *Sd-RanGAP* and *In(2R)Mal*. The functional role of *In(2R)Mal* for drive is still unclear: do these inversions function to suppress recombination between *Sd-RanGAP* and a major distal enhancer on *2R*, or do they contain a major enhancer?

Here, we combine genetic and population genomic approaches to study *SD-Mal* haplotypes sampled from a single population in Zambia, the putative ancestral range of *D. melanogaster* (Pool *et al*. 2012). We address four issues. First, we reveal the structural features of the *SD-Mal* haplotype, including the organization of the insensitive *Rsp* allele and the *In(2R)Mal* rearrangements. Second, we characterize the genetic function of *In(2R)Mal* and its role in drive. Third, we infer the population genetic history of the rapid rise in frequency of *SD-Mal* in Zambia. And fourth, we explore the evolutionary consequences of reduced recombination on *SD-Mal* haplotypes. Our results show that *SD-Mal* experienced a recent chromosome-scale selective sweep mediated by epistatic selection and has, as a consequence of its reduced population recombination rate, accumulated excess non-synonymous mutations and transposable element insertions. The *SD-Mal* haplotype is a supergene that evolves as a small, semi-isolated subpopulation in which complementing *SD-Mal* chromosomes can recombine *inter se* via crossing over and with wildtype chromosomes only via gene conversion. These results have implications for supergene evolution and may explain the enigmatic evolutionary turnover of *SD* haplotypes in different populations around the world.

## Results and discussion

To investigate the evolutionary genomics of *SD-Mal*, we sequenced haploid embryos from nine driving *SD-Mal* haplotypes sampled from a single population in Zambia (Brand *et al*. 2015), the putative ancestral range of *D. melanogaster* (Pool *et al*. 2012). Illumina read depth among samples ranged between ~46-67x, (Supplementary Table S1; BioProject PRJNA649752 in NCBI). Additionally, we obtained ~12x coverage with long-read Nanopore sequencing of one homozygous viable line, *SD-ZI125*, to create a *de novo* assembly of a representative *SD-Mal* haplotype. We use these data to study the evolution of *SD-Mal* structure, diversity, and recombination.

### Chromosomal features of the SD-Mal supergene

The *SD-Mal* haplotype has at least three key features: the main drive locus, the *Sd-RanGAP* duplication on 2*L*; an insensitive *Responder* (*Rsp^i^*) in *2R* heterochromatin; and the paracentric *In(2R)Mal* arrangement on chromosome *2R* (Figure 1). We used our long-read and short-read sequence data for *SD-ZI125* to confirm the structure of the duplication (Fig. 1a) and then validated features in the other *SD-Mal* haplotypes. All *SD-Mal* chromosomes have the *Sd-RanGAP* duplication at the same location as the parent gene on chromosome *2L* (see also Brand *et al*. 2015). The *Rsp* locus, the target of *SD*, corresponds to a block of ~120-bp satellite repeats in *2R* heterochromatin (Fig. 1b; Wu *et al*. 1988). The reference genome, *Iso-1*, has a *Rsp^s^* allele corresponding to a primary *Rsp* locus containing two blocks of tandem *Rsp* repeats—*Rsp-proximal* and *Rsp-major*— with ~1000 copies of the *Rsp* satellite repeat interrupted by transposable elements (Khost *et al*. 2017). A small number of *Rsp* repeats exist outside of the primary *Rsp* locus, although they are not known to be targeted by *SD*. There are three of these additional *Rsp* loci in *Iso-1*: ~10 copies in *2R*, distal to the major *Rsp* locus (*Rsp-minor*); a single copy at the distal end of *2R* (60A); and ~12 copies in *3L* (Houtchens and Lyttle; Larracuente 2014; Khost *et al*. 2017). The genomes of *SD* flies carry ~20 copies of *Rsp* (Wu *et al*. 1988; Pimpinelli and Dimitri 1989), but the organization of the primary *Rsp* locus on *SD* chromosomes is unknown. To characterize the *Rsp* locus of the *SD-Mal* haplotype, we mapped *SD-Mal* reads to an *Iso-1* reference genome (see Khost *et al*. 2017). As expected, reads from *Iso-1* reference are distributed across the whole *Rsp-major* region. For *SD-Mal* chromosomes, however, very few reads map to the *Rsp* repeats at the *Rsp-major* (Fig. 1b). This suggests that all *SD-Mal* have a complete deletion of the primary *Rsp* locus containing *Rsp-proximal* and *Rsp-major* and that the only *Rsp* copies in the *SD-Mal* genomes are the minor *Rsp* loci in chromosomes *2R* and *3L* (Suppl. Fig. S1).

**Figure 1.**
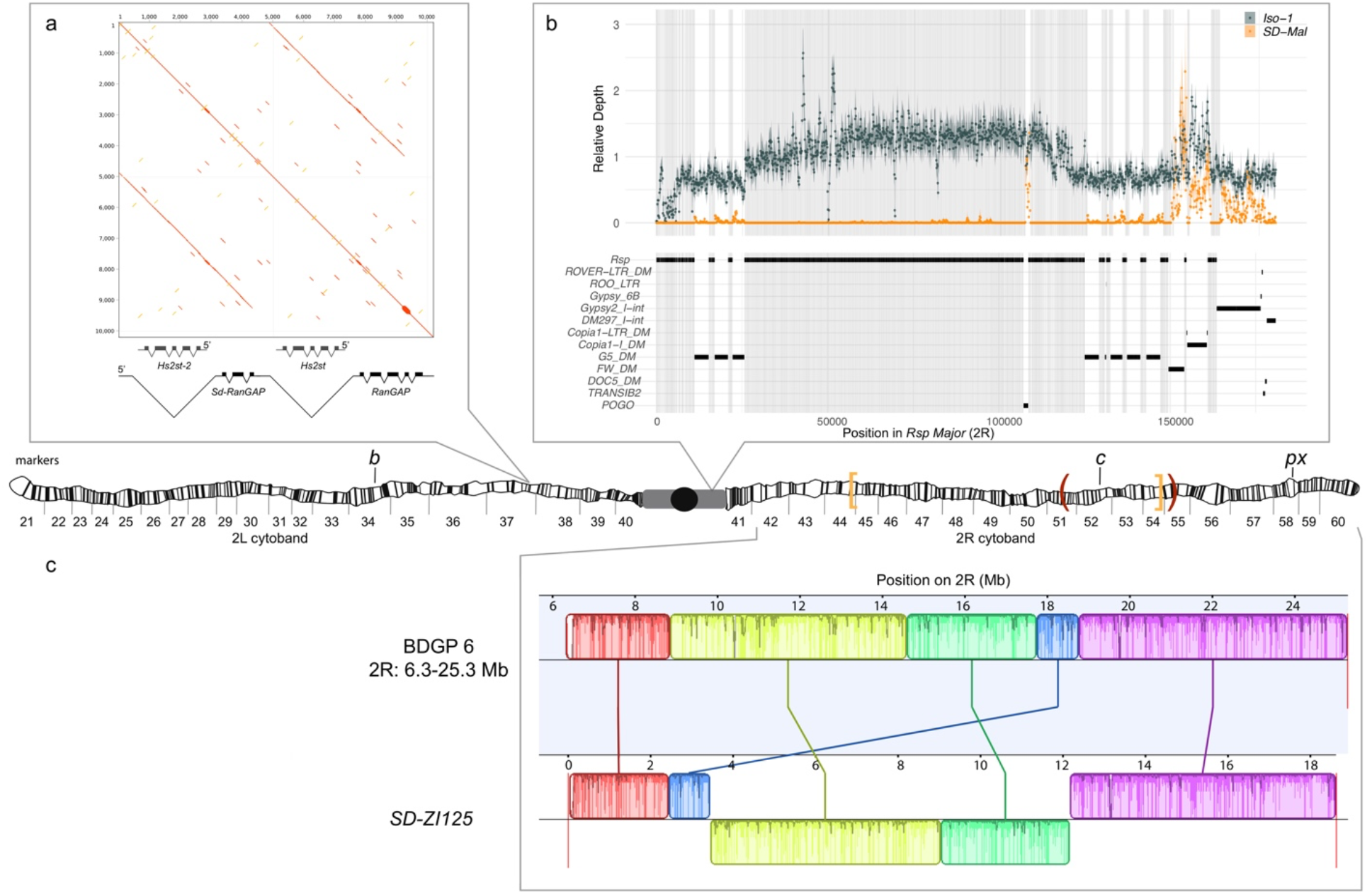
Map depicting the chromosomal features of the *SD-Mal* chromosome. The schematic shows the cytogenetic map of chromosomes *2L* and *2R* (redrawn based on images in (Lefevre 1976)) and the major features of the chromosome. (a) Dotplot showing that the *Sd* locus is a partial duplication of the gene *RanGAP* (in black), located at band *37D2-6*. (b) The *Rsp-major* locus is an array of tandem repeats located in the pericentric heterochromatin (band *h39*). Read mapping shows that *SD-Mal* chromosomes do not have any *Rsp* repeats in the *Rsp-major* locus, consistent with being insensitive to distortion by *Sd* (*Rsp^i^*) (orange, high relative coverage regions correspond to transposable element interspersed), in contrast with *Iso-1*, which is sensitive (*Rsp^s^*). (c) Two paracentric, overlapping inversions constitute the *In(2R)Mal* arrangement: *In(2R)51BC;55E (In(2R)Mal-p)* in orange brackets and *In(2R)44F;54E (In(2R)Mal-d)* in red parentheses). Pericentromeric heterochromatin and the centromere are represented by a grey rectangle and black circle, respectively. Our assembly based on long-read sequencing data provide the exact breakpoints of *In(2R)Mal* and confirms that the distal inversion (*Dmel.r6, 2R:14,591,034-18,774,475*) occurred first, and the proximal inversion (*Dmel.r6, 2R*:8,855,601-15,616,195) followed, overlapping ~1Mb with the distal inversion. The colored rectangles correspond to locally collinear blocks of sequence. Blocks below the center black line indicate regions that align in the reverse complement orientation. Vertical red lines indicate the end of the assembled chromosomes. Visible marker locations used for generating recombinants (*b* (34D1), *c* (52D1), and *px* (58E4-58E8)) are indicated on the cytogenetic map.

The complex *In(2R)Mal* inversion is distal to the *Rsp* locus on chromosome *2R* (Fig 1c). We used our *SD-ZI125* assembly to determine the precise breakpoints of these inversions. Relative to the standard *D. melanogaster 2R* scaffold (BDGP6), *SD-ZI125* has three large, rearranged blocks of sequence corresponding to *In(2R)Mal* (Fig. 1c): a 1.03-Mb block collinear with the reference but shifted proximally; a second inverted 5.74-Mb block; and a third inverted 3.16-Mb block. From this organization, we infer that the distal inversion, which we refer to as *In(2R)Mal-d*, occurred first and spanned 4.18 Mb (approx. *2R*:14,591,003-18,774,475). The proximal inversion, which we refer to as *In(2R)Mal-p*, occurred second and spanned 6.76 Mb, with 1.02 Mb overlapping with the proximal region of *In(2R)Mal-d* (approx. *2R*:8,855,602-17,749,310; see Suppl. Fig. S2). All four breakpoints of the *In(2R)Mal* rearrangement involve simple joins of unique sequence. Three of these four breakpoints span genes: *sns* (*2R*:8,798,489 - 8,856,091), *CG10931* (*2R*:17,748,935 −17,750,136), and *Mctp* (*2R*:18,761,758 - 18,774,824). The CDSs of both *sns* and *Mctp* remain intact in the *In(2R)Mal* arrangement, with the inversion disrupting their 3’UTRs. Neither of these two genes is expressed in testes (https://flybase.org/reports/FBgn0024189; https://flybase.org/reports/FBgn0034389; Chintapalli *et al*. 2007; FB2021_06; Larkin *et al*. 2021), so it is unlikely that they are related to drive. *In(2R)Mal-p* disrupted the CDS of *CG10931*, which is a histone methyltransferase with high expression levels in testis (https://flybase.org/reports/FBgn0034274; Chintapalli *et al*. 2007; FB2021_06; Larkin *et al*. 2021). Future work is required to determine if this gene has a role in the *SD-Mal* drive phenotype.

In African populations, *In(2R)Mal* appears essential for *SD* drive: recombinant chromosomes bearing *Sd* but lacking *In(2R)Mal* do not drive (Presgraves *et al*. 2009; Brand *et al*. 2015). The functional role of *In(2R)Mal* in drive is however unclear. As expected, *In(2R)Mal* suppresses recombination: in crosses between a multiply marked chromosome *2*, *b c px*, and *SD-Mal*, we find that *In(2R)Mal* reduces the *b – c* genetic distance by 54.6% (from 26.6 to 11.8 cM) and the *c* – *px* genetic distance by 92.4% (from 23.2 to 1.8 cM), compared with control crosses between *b c px* and Oregon-R. Our crosses also confirm that *In(2R)Mal* is indeed required for drive: all *Sd, In(2R)Mal* haplotypes show strong drive (Table 1, rows 1 and 2), whereas none of the recombinants that separate *Sd* and *In(2R)Mal* drive (Table 1, rows 3 and 4). We conclude that *SD-Mal* drive requires both *Sd* and *In(2R)Mal*, which implies that one or more essential enhancers, or co-drivers, is located within or distal to *In(2R)Mal*.

**Table 1:** Strength of segregation distortion in recombinants of *SD-ZI125*.

|  | Genotype | Markers | *N* | *n* | +/- *SE* | *k* | +/- *SE* | *k* * | +/- *SE* | *p* -value<br>(*k* * = 0.5) |
| --- | --- | --- | --- | --- | --- | --- | --- | --- | --- | --- |
| 1 | *Sd,In(2R)Mal* | +++; *b*++ | 112 | 90.3 | 6.03 | 0.99 | 0.00 | 0.98 | 0.00 | < 0.0001 |
| 2 | *Sd,In(2R)Mal* | ++*px* | 71 | 118.8 | 9.48 | 0.97 | 0.01 | 0.96 | 0.01 | < 0.0001 |
| 3 | *Sd,In(2R)Mal*<sup>+</sup> | +*c px* | 19 | 147.6 | 14.39 | 0.54 | 0.01 | 0.51 | 0.01 | 0.3082 |
| 4 | *Sd*<sup>+</sup>,*In(2R)Mal* | *b*++ | 24 | 124.8 | 10.31 | 0.68 | 0.02 | 0.55 | 0.03 | 0.0572 |
| 5 | *Sd*<sup>+</sup>,*In(2R)Mal*<sup>+</sup> | +*c px* | 65 | 120.4 | 8.32 | 0.53 | 0.01 | 0.51 | 0.01 | 0.3586 |

Chromosome 2 markers are *black* (*b*), *curved* (*c*), and *plexus* (*px*). *N*, number of crosses; *n*, average number of progeny from the crosses; *SE*, standard error; *k*, average proportion of progeny inheriting the partial *SD<sub>r</sub>* chromosome from *SD<sub>r</sub>/bcpx* males; *k*\*, average proportion of progeny inheriting the partial *SD<sub>r</sub>* chromosome from *SD<sub>r</sub>/bcpx* males, corrected for viability. *P*-values reported by a single sample t-test with a null hypothesis of *k*\*=0.5, as expected for Mendelian segregation.

The temporal order of inversions (first *In(2R)Mal-d*, then *In(2R)Mal-p*) suggests two possible scenarios. *In(2R)Mal-d*, occurring first, may have captured the essential enhancer, with the subsequent *In(2R)Mal-p* serving to further reduce recombination between *Sd* and the enhancer. Alternatively, an essential enhancer is located distal to *In(2R)Mal-d*, and the role of both *In(2R)Mal* inversions is to reduce recombination with *Sd*. To distinguish these possibilities, we measured drive in *b^+^ Sd c^+^ In(2R)Mal px* recombinants, which bear *Sd* and *In(2R)Mal* but recombine between the distal breakpoint of *In(2R)Mal (2R*: 18,774,475) and *px (2R: 22,494,297)*. All of these recombinants show strong drive (n=71; Table 1, row 2). Assuming that recombination is uniformly distributed throughout the 3.72-Mb interval between the *In(2R)Mal-d* distal breakpoint and *px*, the probability of failing to separate an essential codriver or distal enhancer among any of our 71 recombinants is <0.014. Furthermore, using molecular markers, we detected two recombinants within 100 kb of the distal breakpoint of *In(2R)Mal*, both with strong drive (*k*>0.99). We therefore infer that the co-driver likely resides within the *In(2R)Mal* arrangement. More specifically, we speculate that the *In(2R)Mal-d* inversion both captured the co-driver and reduced recombination with *Sd*, whereas *In(2R)Mal-p* tightened linkage between centromere-proximal components of *SD-Mal* and *In(2R)Mal-d*.

While recombination occurs readily between *Sd* and the proximal break of *In(2R)Mal* (Table 1; Presgraves *et al*. 2009; Brand *et al*. 2015), long-range linkage disequilibrium nevertheless exists between *Sd* and *In(2R)Mal*. Using 204 haploid genomes from Zambia (Lack *et al*. 2016; see Methods), we identified 198 wildtype haplotypes (*Sd*^+^ *In(2R)Mal^+^*), 3 *SD-Mal* haplotypes (*Sd In(2R)Mal*), 3 recombinant haplotypes (all *Sd In(2R)Mal^+^*, none *Sd^+^ In(2R)Mal*). Despite the individually low frequencies of *Sd* (frequency = 0.0294) and *In(2R)Mal* (frequency = 0.0147), they tend to co-occur on the same haplotypes (Fisher’s exact *P*=1.4 × 10^−5^). Given a conservative recombination frequency between *Sd* and *In(2R)Mal* of ~5 cM (FlyBase; FB2021_06; Larkin *et al*. 2021), the observed estimated coefficient of linkage disequilibrium, *D* = 0.0143, has a half-life of just ~14 generations (1.4 years, assuming 10 generations per year) and should decay to negligible within 100 generations (10 years). We therefore conclude that strong epistatic selection maintains the *SD-Mal* supergene haplotype.

### Rapid increase in frequency of the SD-Mal supergene

We used population genomics to infer the evolutionary history and dynamics of *SD-Mal* chromosomes. We called SNPs in our Illumina reads from nine complete *SD-Mal* haplotypes from Zambia (see Methods). For comparison, we also analyzed wildtype (*SD^+^*) chromosomes from the same population in Zambia (Lack *et al*. 2016), including those with chromosome *2* inversions: 10 with the *In(2L)t* inversion and 10 with the *In(2R)NS* inversion (see Methods). Table 2 shows that nucleotide diversity (*π*) is significantly lower on *SD-Mal* haplotypes compared to uninverted *SD*^+^ chromosome arms (Table 2, rows 1 and 4; Figure 2a). The relative reduction in diversity on *SD-Mal* haplotypes is distributed heterogeneously: *π* is sharply reduced for a large region that spans ~25.8 Mb, representing 53% of chromosome 2 and extending from *Sd-RanGAP* on *2L (2L*: 19,441,959; Suppl. Fig. S3), across the centromere, and to ~2.9 Mb beyond the distal breakpoint of *In(2R)Mal (2R*: 18,774,475; Table 2, rows 3, 5 and 6; Fig. 2a). Thus, the region of reduced nucleotide diversity on *SD-Mal* chromosomes covers all of the known essential loci for the drive phenotype: *Sd-RanGAP, Rsp^i^* and *In(2R)Mal*.

**Table 2:**
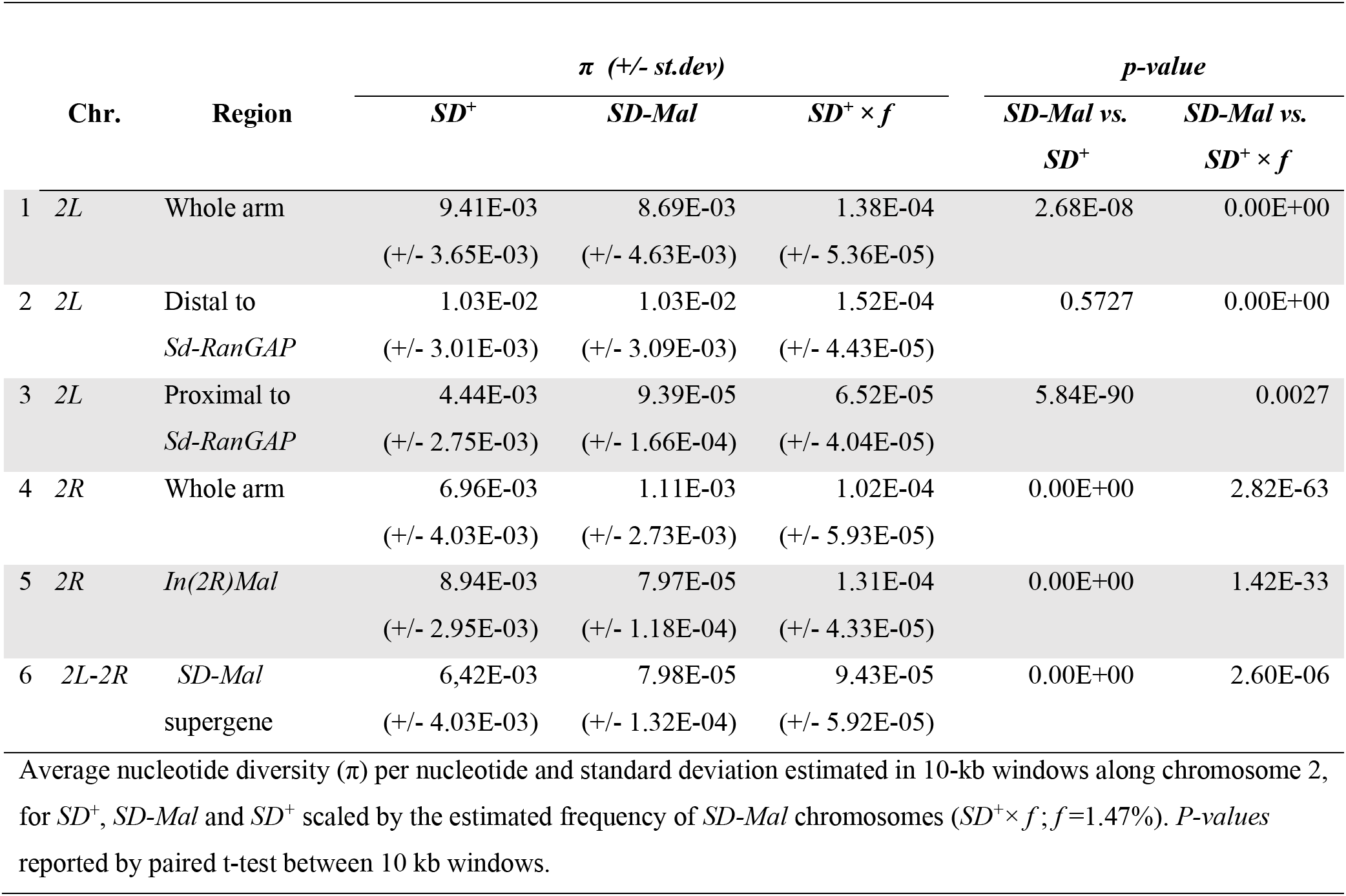
Nucleotide diversity (π) on *SD-Mal* an *SD*^+^ chromosomes.

**Figure 2.**
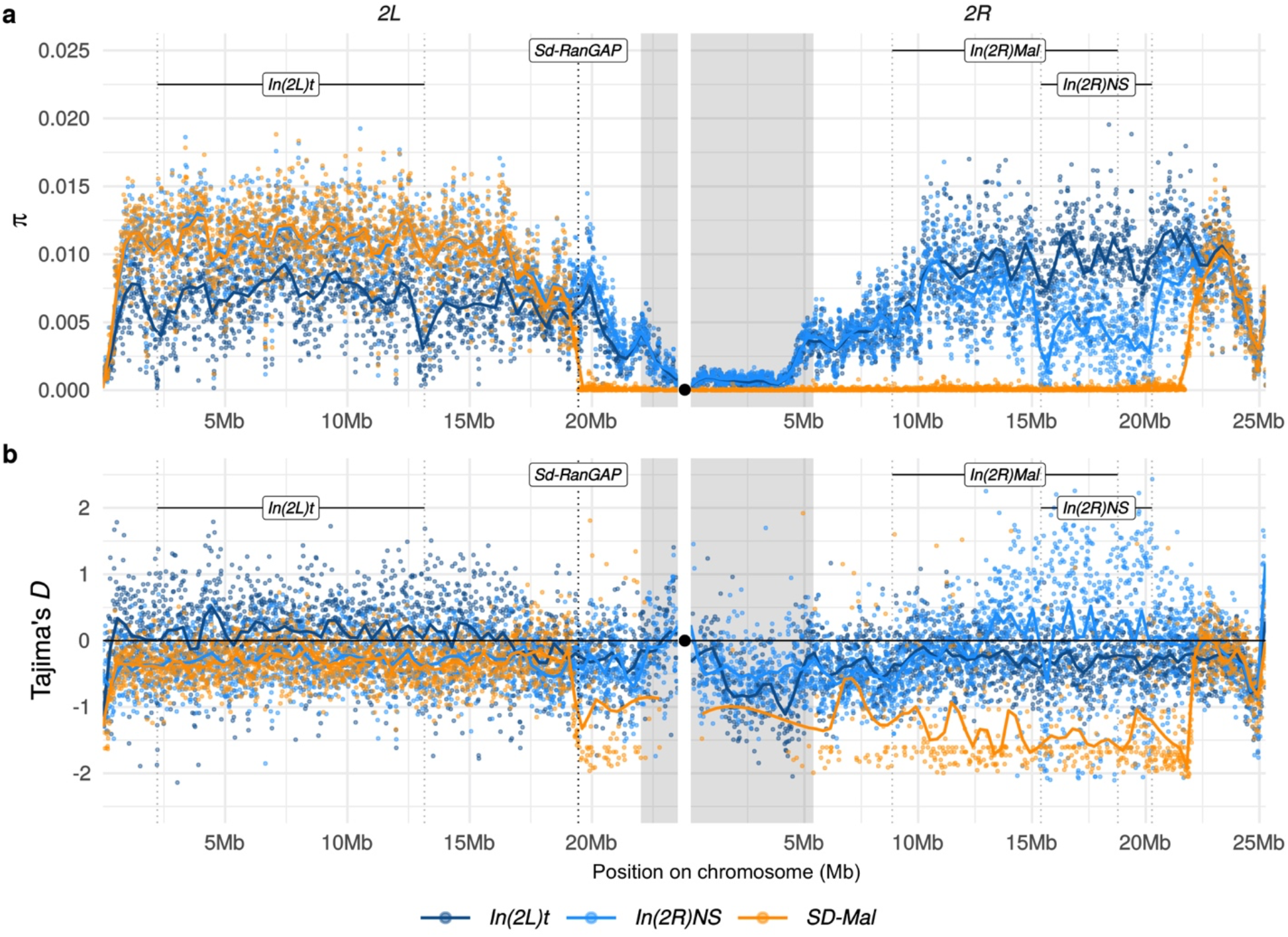
Diversity on *SD-Mal* chromosomes. (a) Average pairwise nucleotide diversity per site (π) and (b) Tajima’s *D* estimates in non-overlapping 10-kb windows along chromosome *2*, in Zambian *SD-Mal* chromosomes (n=9, orange) and *SD*^+^ chromosomes from the same population, bearing the cosmopolitan inversions *In(2L)t* (n=10, dark blue) and *In(2R)NS* (n=10, light blue). Regions corresponding to pericentric heterochromatin are shaded in grey and the centromere location is marked with a black circle. *SD-Mal* chromosomes show a sharp decrease in nucleotide diversity and skewed frequency spectrum from the *Sd* locus (*Sd-RanGAP, 2L*:19.4Mb) to ~2.9 Mb beyond the distal breakpoint of *In(2R)Mal*.

The reduced nucleotide diversity among *SD-Mal* may not be surprising, given its low frequency in natural populations (see below; Presgraves *et al*. 2009; Brand *et al*. 2015). *SD* persists at low frequencies in populations worldwide, presumably reflecting the balance between drive, negative selection, and genetic suppression and/or resistance (Hartl 1975; Charlesworth and Hartl 1978; Larracuente and Presgraves 2012). If the *SD-Mal* supergene has been maintained at stable drive-selection-suppression equilibrium frequency for a long period of time, then its neutral nucleotide diversity may reflect a mutation-drift equilibrium appropriate for its effective population size. Under this scenario, we expect diversity at the supergene to be similar to wild type (*SD^+^*) diversity scaled by the long-term equilibrium frequency of *SD*. We estimated *SD-Mal* frequency to be 1.47% by identifying the *Sd* duplication and *In(2R)Mal* breakpoints in 204 haploid genomes from Zambia (3/204, comparable to Presgraves *et al*. 2009; Brand *et al*. 2015; data from Lack *et al*. 2016; see Methods). To approximate our expectation under mutation-drift equilibrium, we scaled average *π* from the *SD*^+^ sample by 1.47% in 10-kb windows across the region corresponding to the *SD-Mal* supergene, defined as the region from *Sd-RanGAP* to the distal breakpoint of *In(2R)Mal*. Table 2 (row 6) shows that diversity in the *SD-Mal* supergene region is still significantly lower than expected, suggesting that the low frequency of *SD-Mal* cannot fully explain its reduced diversity. This observation suggests two possibilities: the *SD-Mal* supergene historically had an equilibrium frequency less than 1.47% in Zambia; or the *SD-Mal* supergene, having reduced recombination, has experienced hitchhiking effects due to background selection and/or a recent selective sweep.

To distinguish between these possibilities, we analyzed summaries of the site frequency spectrum. We find strongly negative Tajima’s *D* mirroring the distribution of reduced diversity, indicating an excess of rare alleles (Figure 2b). Such a skew in the site frequency spectrum suggests a recent increase in frequency of the *SD-Mal* supergene in Zambia. The high differentiation of *SD-Mal* from *SD*^+^ chromosomes from the same population similarly suggests a large shift in allele frequencies. Wright’s fixation index, *F_ST_*, in the *SD-Mal* supergene region is unusually high for chromosomes from the same population (Figure 3a). Neither of the *SD*^+^ chromosomes with cosmopolitan inversions show such high differentiation within their inversions, and mean nucleotide differences (*d_XY_*) between *SD-Mal* and *SD*^+^ is comparable to the other inversions, implying that the differentiation of the *SD-Mal* supergene is recent. Our results are thus consistent with a rapid increase in frequency of the *SD-Mal* haplotype that reduced nucleotide diversity within *SD-Mal* and generated large differences in allele frequencies with *SD*^+^ chromosomes.

**Figure 3.**
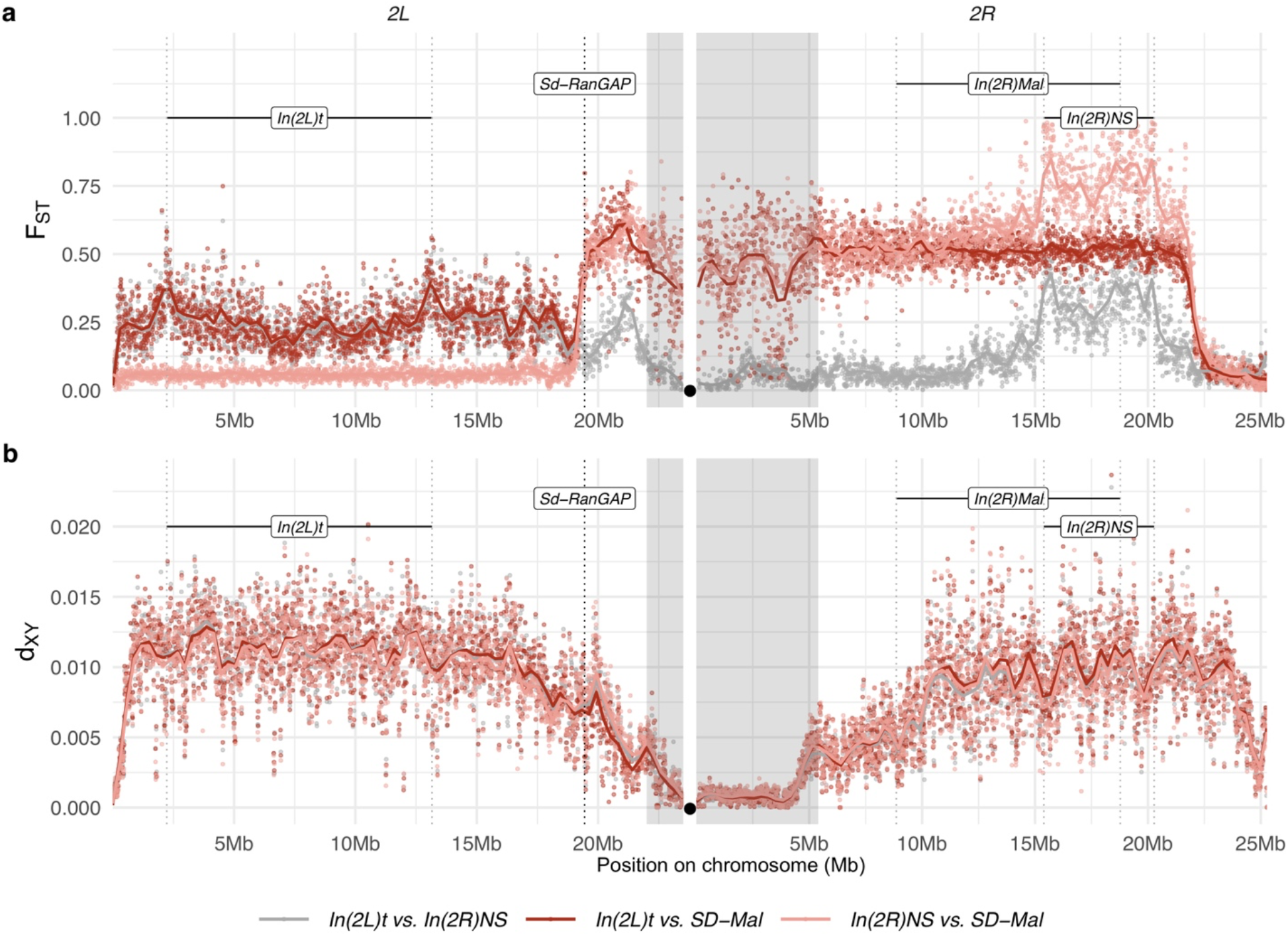
Differentiation between *SD-Mal* and wildtype chromosomes. (a) Pairwise *F_ST_* and (b) *d_XY_* per base pair in non-overlapping 10-kb windows along chromosome *2*, between Zambian *SD-Mal* haplotypes (n=9) and wildtype chromosomes from the same population, bearing the cosmopolitan inversions *In(2L)t* (n=10) and *In(2R)NS* (n=10). Regions corresponding to pericentric heterochromatin are shaded in grey and the centromere location is marked with a black circle.

To estimate the timing of the recent expansion of the *SD-Mal* supergene, we used an Approximate Bayesian Computation (ABC) method with rejection sampling in neutral coalescent simulations. We do not know if *SD* chromosomes acquired *In(2R)Mal* in Zambia or if the inversions occurred *de novo* on an *SD* background. For our simulations, we assume that the acquisition of these inversion(s) was a unique event that enhanced drive strength and/or efficiency. Under this scenario, extant *SD-Mal* chromosomes have a single origin. We therefore simulated this history in a coalescent framework as an absolute bottleneck to a single chromosome. We performed simulations considering a sample size of *n* = 9 and assumed no recombination in the ~9.92-Mb region of *In(2R)Mal*. We simulated with values of *S* drawn from a uniform distribution ±5% of the observed number of segregating sites in *In(2R)Mal*. We considered a prior uniform distribution of the time of the expansion (*t*) ranging from 0 to *4Ne* generations (0 – 185,836 years ago), assuming that *D. melanogaster N_e_* in Zambia 3,160,475 (Kapopoulou *et al*. 2018), a *In(2R)Mal* frequency of 1.47%, and 10 generations per year (Li and Stephan 2006; Thornton and Andolfatto 2006; Laurent *et al*. 2011; Kapopoulou *et al*. 2018). Using the ABC with rejection sampling conditional on our observed estimates of *π* and Tajima’s *D* for *In(2R)Mal* (*π_In(2R)Mal_* = 760.49, *D* = −1.27; note that *π_In(2R)Mal_* is an overall, unscaled estimate of nucleotide diversity for the whole *In(2R)Mal* region), we infer that the *SD-Mal* expansion began ~0.096 (95% credibility intervals 0.092 - 0.115) × 4*N_e_* generations ago or, equivalently, ~1792 years ago (0.88% rejection sampling acceptance rate; Figure 4). To account for possible gene conversion (see below), we discarded SNPs shared with *SD*^+^ chromosomes (see below), and recalculated *π* and Tajima’s *D* using only private SNPs (*π_In(2R)Mal_* = 563.34, *D* = −1.41). Based on these parameters, the estimated *SD-Mal* expansion occurred ~0.0737 (95% credibility intervals 0.070 - 0.092) 4*N_e_* generations ago, ~1368 years (0.86% rejection sampling acceptance rate Figure 4). To calculate the posterior probability of the model, we performed 100,000 simulations under a model assuming a stable frequency of *SD-Mal* and under sweep models (assuming *t*_all_ = 0.096 and *t*_shared_excl_ 0.0737) (Suppl. Fig. S4). The simulated data are inconsistent with a long-term stable frequency of *SD-Mal* (All SNPs, p_*π*_ = 0.0526, p_*D*_ = 0.1084; Private, p_*π*_ = 0.0285, p*_D_* = 0.0755). Instead, our simulations suggest that a recent selective sweep is more consistent with the data (All SNPs, p_*π*_ = 0.3106, p*_D_* = 0.5929; Private, p_*D*_ = 0.3091, p*_D_* = 0.6092). Taken together, evidence from nucleotide diversity, the site frequency spectrum, population differentiation, and coalescent simulations suggest a rapid non-neutral increase in frequency of the *SD-Mal* supergene that began < 2,000 years ago.

**Figure 4.**
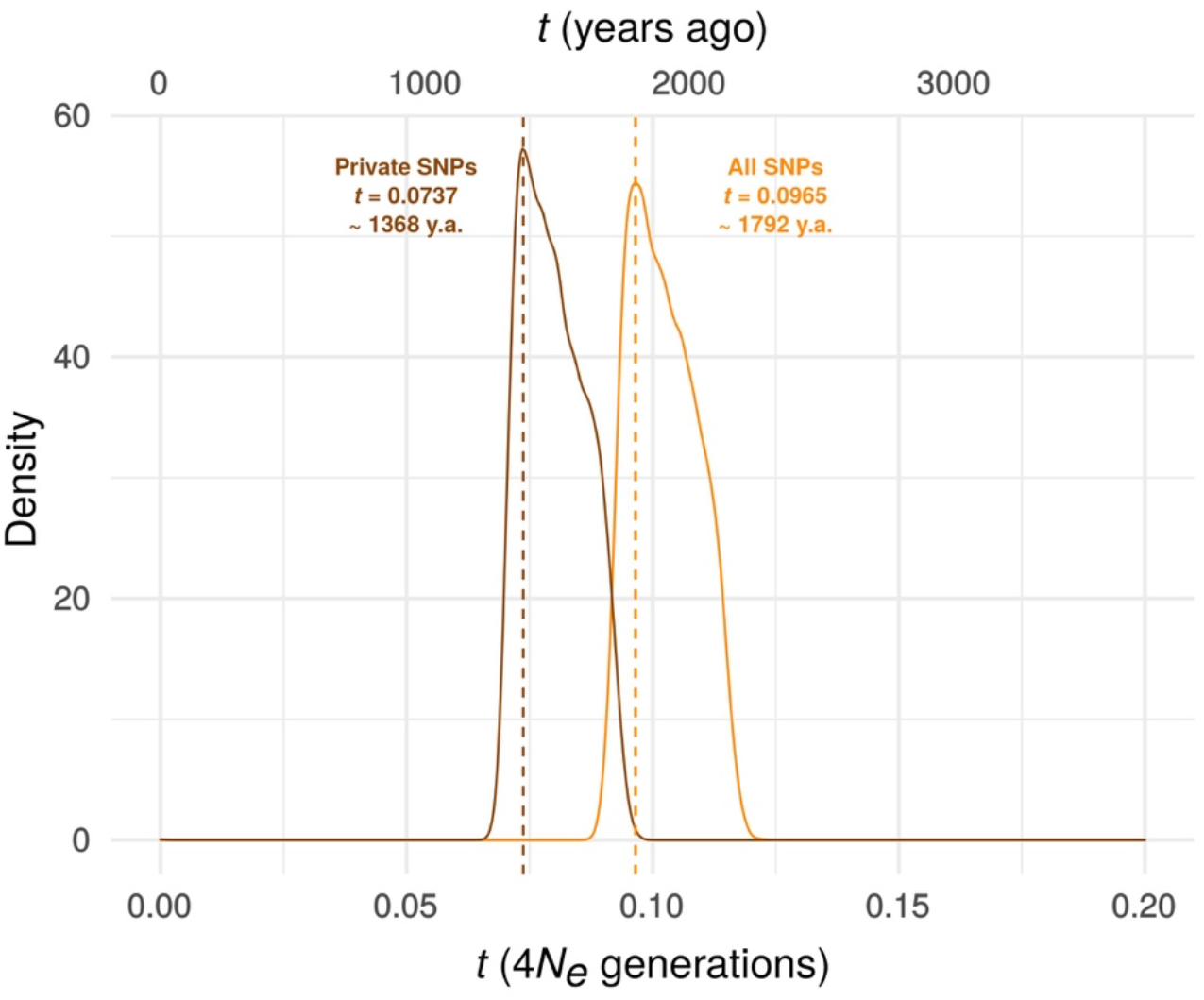
Estimating the time since the *SD-Mal* selective sweep. ABC estimates based on 10,000 posterior samples place the onset of the selective sweep between 0.096 (95% C.I. 0.092 - 0.115) and 0.0737 (0.070 - 0.092) × 4*N_e_* generations, i.e. ~1,368-1,792 years ago, considering recent estimates of *N_e_* in Zambia from (Kapopoulou *et al*. 2018), frequency of *SD-Mal* in Zambia 1.47% and 10 generations per year). Estimates were done considering only *In(2R)Mal*, where crossing over is rare and only occurs between *SD-Mal* chromosomes using all SNPs and excluding shared SNPs in order to account for gene conversion from *SD*^+^ chromosomes.

The sweep signal on the *SD-Mal* haplotypes begins immediately distal to *Sd-RanGAP* on *2L* and extends ~3 Mb beyond the distal boundary of *In(2R)Mal* on *2R*. To understand why the sweep extends so far beyond the *In(2R)Mal-d* distal breakpoint, we consider three, not mutually exclusive, possibilities. First, chromosomal inversions can suppress recombination ~1-3 Mb beyond their breakpoints (Stevison *et al*. 2011; Miller *et al*. 2016; Crown *et al*. 2018; Miller *et al*. 2018), extending the size of the sweep signal. To determine the extent of recombination suppression caused by *In(2R)Mal*, we estimated recombination rates in the region distal to the inversion. The expected genetic distance between the distal breakpoint of *In(2R)Mal (2R*: 18.77 Mb) and *px (2R*: 22.49 Mb) is ~13.87 cM (Fiston-Lavier *et al*. 2010). Measuring recombination between *SD-Mal* and standard arrangement chromosomes for the same (collinear) interval, we estimate a genetic distance of ~1.76 (see above), an 87.3% reduction. *In(2R)Mal* strongly reduces recombination beyond its bounds. Second, although we have inferred that the essential enhancer resides within the *In(2R)Mal* inversion (see above), we have not excluded the possibility of weak enhancers distal to the inversion which might contribute to the sweep signal. We find that *SD-Mal* chromosomes with *In(2R)Mal-distal* material recombined away (*b^+^ Sd c^+^ In(2R)Mal px*) have modestly but significantly lower drive strength (*k* = 0.96 *versus* 0.98; Table 1, lines 1-2), suggestive of one or more weak distal enhancers. Third, there may be mutations distal to *In(2R)Mal* that contribute to the fitness of *SD-Mal* haplotypes but without increasing the strength of drive, *e.g*., mutations that compensate for the effects of *SD-Mal*-linked deleterious mutations.

### Recombination on SD supergenes

While nearly all *SD-Mal* haplotypes are individually homozygous lethal and do not recombine with wild type chromosomes in and around *In(2R)Mal*, ~90% of pairwise combinations of different *SD-Mal* chromosomes (*SD_i_/SD_j_*) are viable and fertile in complementation tests (Presgraves *et al*. 2009; Brand *et al*. 2015). Therefore, recombination may occur between *SD-Mal* chromosomes in *SD_i_/SD_j_* heterozygous females. To determine if *SD-Mal* chromosomes recombine, we estimated mean pairwise linkage disequilibrium (*r^2^*) between SNPs located within the *In(2R)Mal* arrangement. We found that mean *r^2^* between pairs of SNPs declines as a function of the physical distance separating them (Figure 5a), a hallmark of recombination via crossing over (Hill and Robertson 1968; Miyashita and Langley 1988; Schaeffer and Miller 1993; Awadalla *et al*. 1999; Conway *et al*. 1999). Pairwise LD is higher and extends further in *In(2R)Mal* than in the equivalent region of *SD*^+^ chromosomes or in any of the other two cosmopolitan inversions, *In(2L)t* and *In(2R)NS* (Figure 5a). This pattern is not surprising: the low frequency of *SD-Mal* makes *SD_i_/SD_j_* genotypes, and hence the opportunity for recombination, rare. To further characterize the history of recombination between *SD-Mal* haplotypes, we used 338 non-singleton, biallelic SNPs in *In(2R)Mal* to trace historical crossover events. From these SNPs, we estimate that Rm (Hudson and Kaplan 1985), the minimum number of recombination events, in this sample of *SD-Mal* haplotypes is 15 (Figure 5c). Thus, assuming that these *SD-Mal* haplotypes are ~17,929 generations old (Figure 4), we estimate that recombination events between *SD-Mal* haplotypes occur a minimum of once every ~1,195 generations. We can thus confirm that crossover events are relatively rare, likely due to the low population frequency of *SD-Mal* and the possibly reduced fitness of *SD_i_/SD_j_* genotypes.

**Figure 5.**
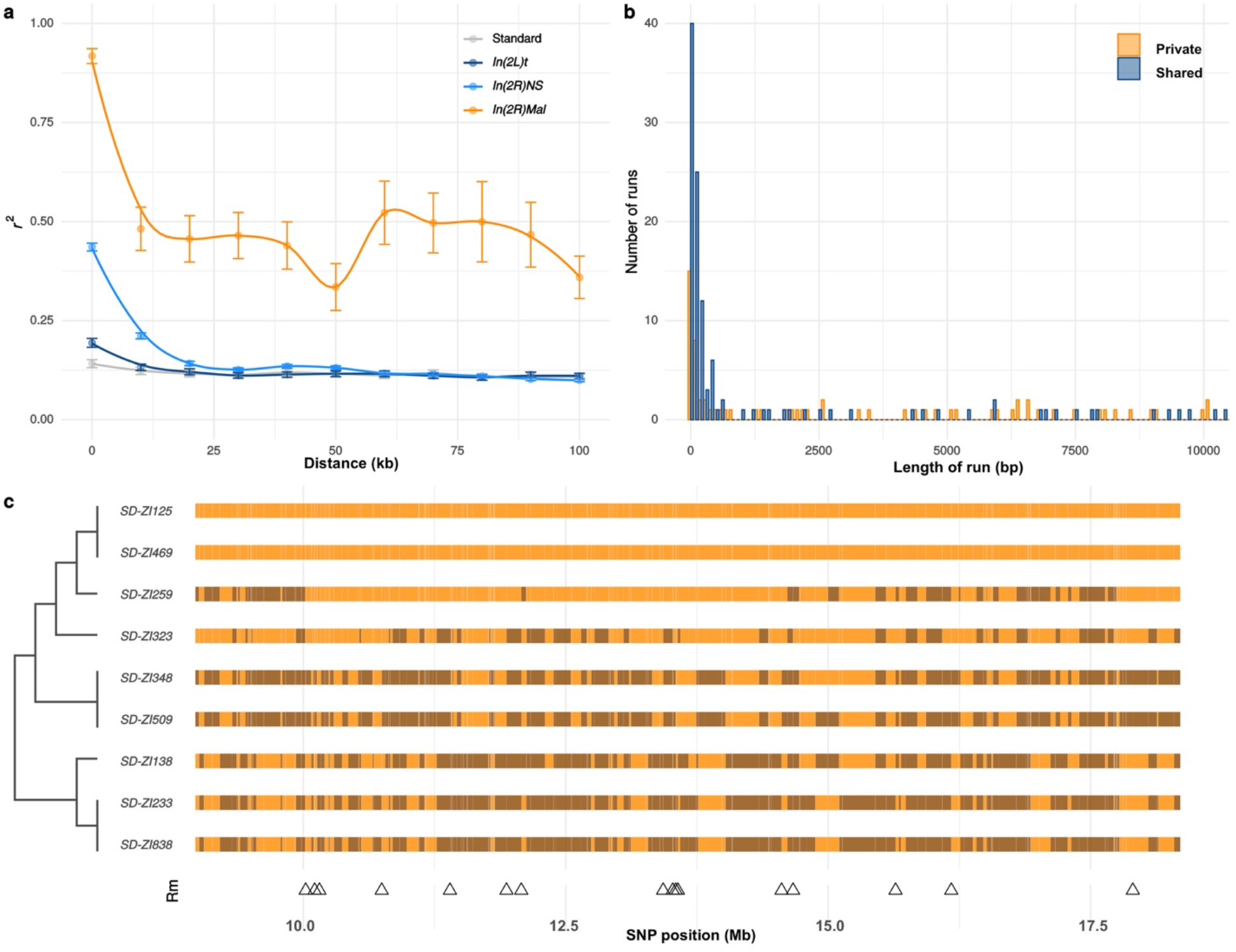
Recombination on *SD-Mal* haplotypes. (a) Linkage disequilibrium (*r^2^*) as a function of distance in 10-kb windows, measured in *In(2R)Mal, In(2L)t, In(2R)NS*, and the corresponding region of *In(2R)Mal* in a standard, uninverted *2R* chromosome. (b) Histogram of length of runs of SNPs in *In(2R)Mal* shows that a high proportion of shared SNPs concentrate in runs shorter than 1 kb. (c) Chromosomal configuration of the 338 non-singleton SNPs in nine different *SD-Mal* lines. Color coded for two states (same in light orange or different in dark orange) using *SD-ZI125* as reference. Locations of minimal number of recombination events are labeled as triangles at the bottom. Maximum likelihood tree is displayed on the left.

While crossing over is suppressed in *SD-Mal/SD*^+^ heterozygotes, gene conversion or double crossover events may still occur, accounting for the shared SNPs between *SD-Mal* and *SD*^+^ chromosomes within *In(2R)Mal*. As both events exchange tracts of sequence, we expect shared SNPs to occur in runs of sites at higher densities than private SNPs, which should be distributed randomly. Accordingly, in *In(2R)Mal*, SNP density is five times higher for runs of shared SNPs (0.63 SNPs/kb) than for runs of *SD*-private SNPs (0.12 SNPs/kb), as expected if *SD*^+^ chromosomes, which have higher SNP densities, were donors of conversion tract sequences. Additionally, 62.2% (89 out of 143) of the shared SNP runs are <1 kb (Figure 5b), consistent with current estimates of gene conversion tract lengths in *D. melanogaster* (Comeron *et al*. 2012). Surprisingly, these inferred gene conversion events are unevenly distributed across *In(2R)Mal*, being more frequent in the *In(2R)Mal-p* than in *In(2R)Mal-d* (Suppl. Table S2). Our discovery that *SD-Mal* haplotypes can recombine with each other distinguishes the *SD-Mal* supergene from other completely genetically isolated supergenes (Wang *et al*. 2013; Charlesworth 2016; Tuttle *et al*. 2016). The lack of crossing over with *SD*^+^ chromosomes, however, means that *SD-Mal* haplotypes evolve as a semi-isolated subpopulation, with a nearly 100-fold smaller *N_e_* and limited gene flow from *SD*^+^ via gene conversion events. The reduced recombination, low *N_e_*, and history of epistatic selection may nevertheless lead to a higher genetic load on *SD-Mal* than *SD*^+^ chromosomes. We therefore examined the accumulation of deleterious mutations, including non-synonymous mutations and transposable elements, on the *SD-Mal* supergene.

### Consequences of reduced recombination, small effective size, and epistatic selection

We first studied the effects of a reduced efficacy of selection on SNPs in *In(2R)Mal*. As many or most non-synonymous polymorphisms are slightly deleterious (Ohta 1976; Fay *et al*. 2001; Eyre-Walker *et al*. 2002), relatively elevated ratios of non-synonymous to synonymous polymorphisms (*N/S* ratio) can indicate a reduced efficacy of negative selection. For the SNPs in *In(2R)Mal*, the overall *N/S* ratio is 2.3-fold higher than that for the same region of *SD*^+^ chromosomes (Table 3). Notably, the *N/S* ratio for private SNPs is 3.1-fold higher (Table 3), whereas the N/S ratios for shared SNPs do not significantly differ from *SD*^+^ chromosomes (Table 3, Suppl. Fig. S5). These findings suggest that gene conversion from *SD*^+^ ameliorates the accumulation of potentially deleterious non-synonymous mutations on *SD-Mal* chromosomes.

**Table 3.**
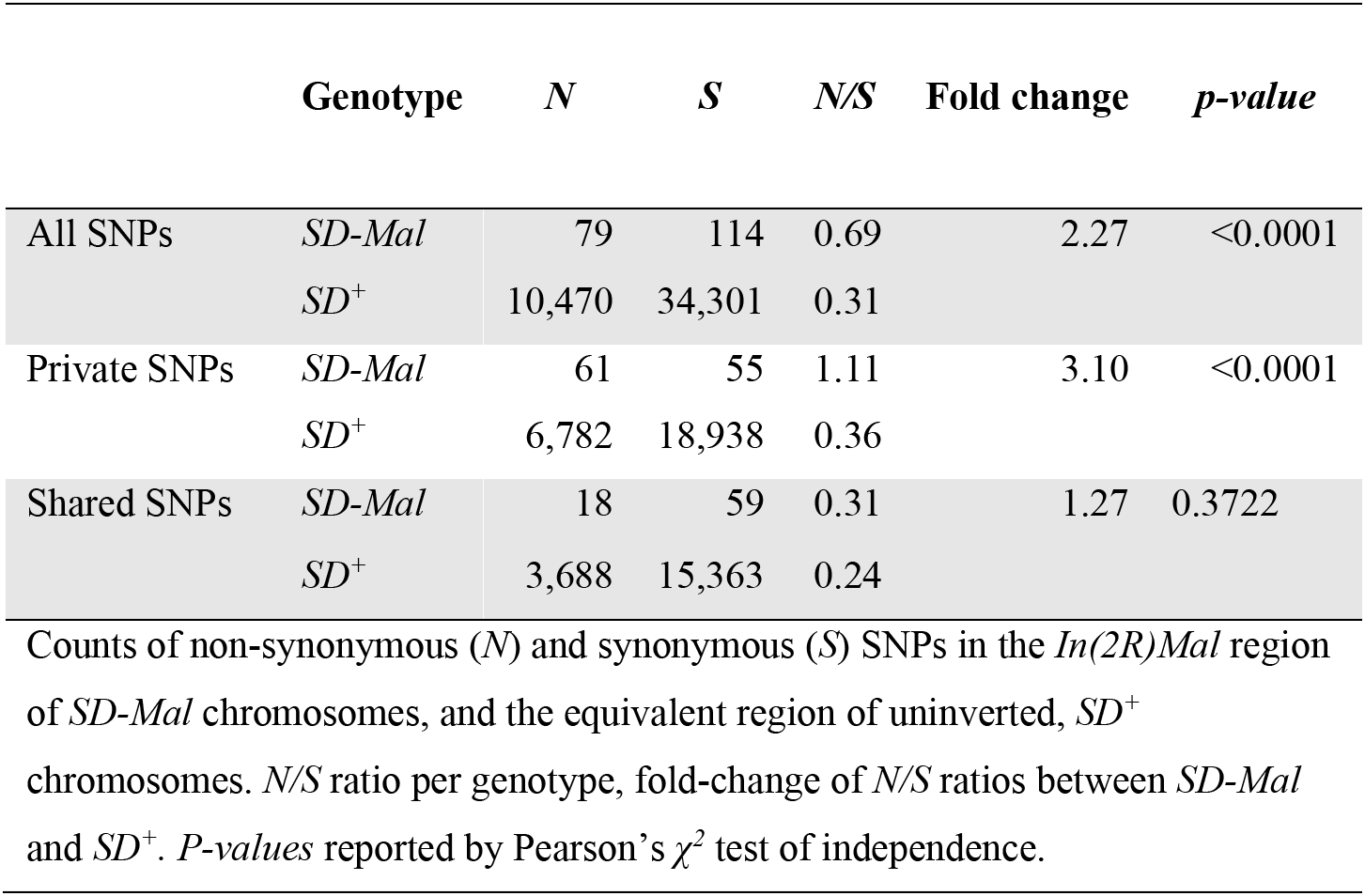
Synonymous and non-synonymous SNPs

|  | Genotype | *N* | *S* | *N/S* | Fold change | *p-value* |
| --- | --- | --- | --- | --- | --- | --- |
| All SNPs | *SD-Mal* | 79 | 114 | 0.69 | 2.27 | <0.0001 |
|  | *SD<sup>+</sup>* | 10,470 | 34,301 | 0.31 |  |  |
| Private SNPs | *SD-Mal* | 61 | 55 | 1.11 | 3.10 | <0.0001 |
|  | *SD<sup>+</sup>* | 6,782 | 18,938 | 0.36 |  |  |
| Shared SNPs | *SD-Mal* | 18 | 59 | 0.31 | 1.27 | 0.3722 |
|  | *SD<sup>+</sup>* | 3,688 | 15,363 | 0.24 |  |  |

Counts of non-synonymous (*N*) and synonymous (*S*) SNPs in the *In(2R)Mal* region of *SD-Mal* chromosomes, and the equivalent region of uninverted, *SD<sup>+</sup>* chromosomes. *N/S* ratio per genotype, fold-change of *N/S* ratios between *SD-Mal* and *SD<sup>+</sup>*. *P-values* reported by Pearson's $\chi^2$ test of independence.

Gene conversion may not, however, rescue *SD-Mal* from deleterious transposable elements (TE) insertions, as average TE length exceeds the average gene conversion tract length (Kaminker *et al*. 2002). TEs accumulate in regions of reduced recombination, such as centromeres (Charlesworth *et al*. 1994) and inversions, especially those at low frequency (Eanes *et al*. 1992; Sniegowski and Charlesworth 1994). Indeed, TE densities for the whole euchromatic region of chromosome *2R* are significantly higher for *SD-Mal* compared to *SD*^+^ chromosomes (Figure 6a). This increased TE density on *SD-Mal* is driven by the non-recombining regions of the haplotype: *In(2R)Mal* has significantly higher TE density than *SD*^+^ whereas the distal region of *2R* outside of the sweep region, does not (Figure 6b). The most overrepresented families in *In(2R)Mal* relative to standard chromosomes are *M4DM, MDG1, ROO_I*, and *LINE* elements (Suppl. Fig.S6)— TEs that are currently or recently active (Kaminker *et al*. 2002; Kofler *et al*. 2015; DÍaz-GonzÁlez and DomÍnguez 2020)— consistent with the recent origin of the *SD-Mal* haplotype. Thus, the differences in shared *versus* private SNPs suggests that gene conversion from *SD*^+^ chromosomes may provide a mechanism to purge deleterious point mutations but not TEs. Despite occasional recombination, the small *N_e_* of *SD-Mal* haplotypes has incurred a higher genetic load.

**Figure 6.**
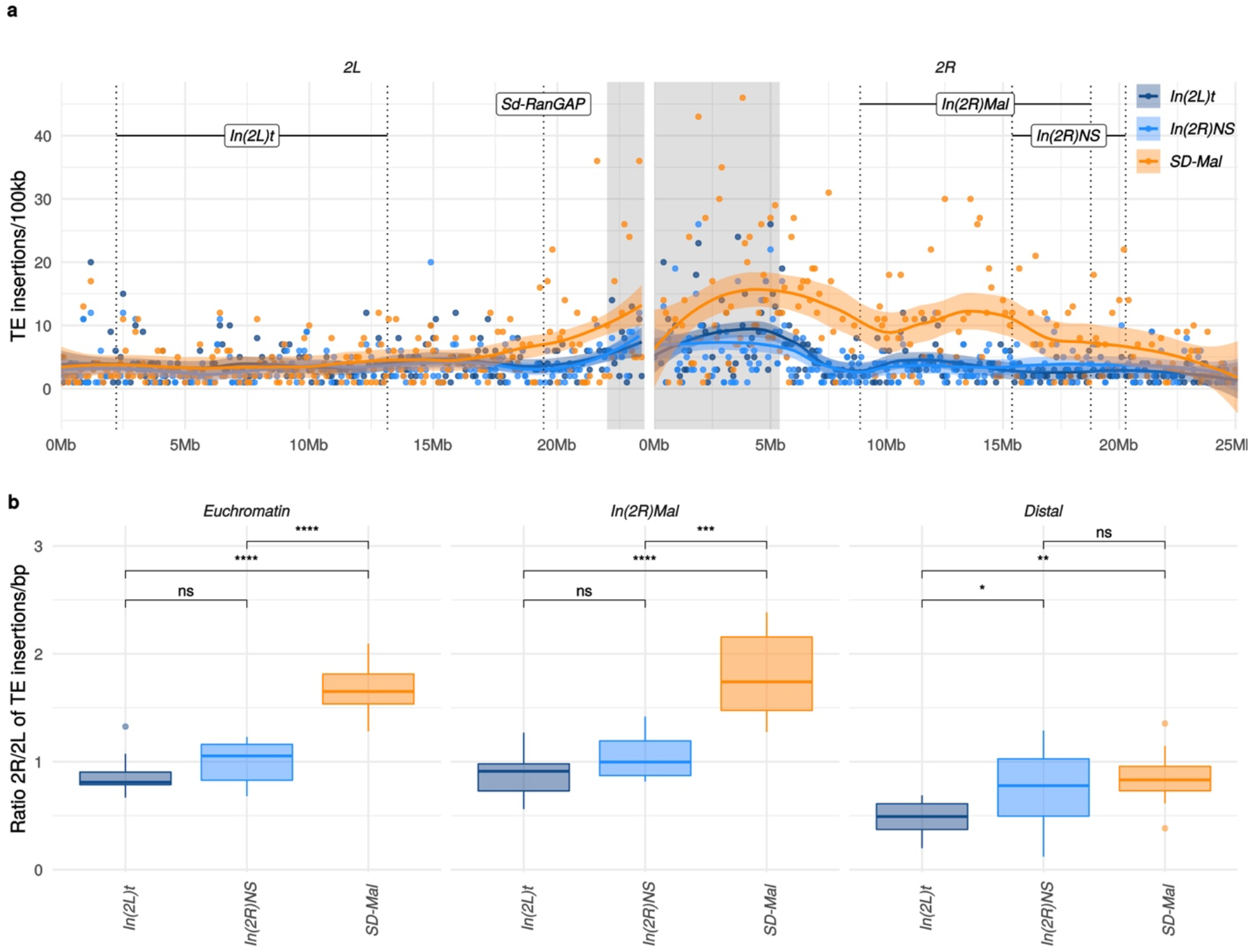
Transposable elements on *SD-Mal* haplotypes. (a) Number of *TE* insertions per 100-kb windows along chromosome 2, in Zambian *SD* chromosomes (n=9, orange) and wildtype chromosomes from the same population, bearing the cosmopolitan inversions *In(2L)t* (n=10, dark blue) and *In(2R)NS* (n=10, light blue). (b) Ratio of the number of insertions in the euchromatin of *2R* to *2L* per library. The relative enrichment in *TE*s in *2R* of *SD-Mal* haplotypes is mostly due to an increase of TE insertions in non-recombining regions of the chromosome.

## Conclusions

Supergenes are balanced, multigenic polymorphisms in which epistatic selection among component loci favors the recruitment of recombination modifiers that reinforce the linkage of beneficial allelic combinations. The advantages of reduced recombination among strongly selected loci can however compromise the efficacy of selection at linked sites. Supergenes thus provide opportunities to study the interaction of recombination and natural selection. We have studied a population of *selfish* supergenes, the *SD-Mal* haplotypes of Zambia, to investigate the interplay of recombination, selection, and meiotic drive. Our findings demonstrate, first, that the *SD-Mal* supergene extends across ~25.8 Mb of *D. melanogaster* chromosome *2*, a region that comprises the driving *Sd-RanGAP*, a drive-insensitive deletion at the major *Rsp* locus, and the *In(2R)Mal* double inversion. Second, using genetic manipulation, we show that *SD-Mal* requires *Sd-RanGAP* and an essential co-driver that localizes almost certainly within the *In(2R)Mal* rearrangement, and probably within the distal inversion. These data provide experimental evidence for epistasis between *Sd-RanGAP* and *In(2R)Mal*: neither allele can drive without the other. Third, we provide population genomics evidence that epistatic selection on loci spanning the *SD-Mal* supergene region drove a very recent, chromosome-scale selective sweep. These patterns are consistent with recurrent episodes of replacement of one *SD* haplotype by others (Presgraves *et al*. 2009; Brand *et al*. 2015). Fourth, despite rare crossovers among complementing *SD-Mal* haplotypes and gene conversion from wildtype chromosomes, the relative genetic isolation and low frequency of *SD-Mal* results in the accumulation of deleterious mutations including, especially, TE insertions. From these findings, we conclude that the *SD-Mal* supergene population is of small effective size, semi-isolated by from the greater population of wildtype chromosomes, and subject to bouts of very strong selection.

Non-recombining supergenes that exist exclusively in heterozygous state tend to degenerate, as in the case of Y chromosomes (reviewed in Charlesworth and Charlesworth 2000) and some autosomal supergenes which, for different reasons, lack any opportunity for recombination (Uyenoyama 2005; Wang *et al*. 2013; Tuttle *et al*. 2016; Branco *et al*. 2018; Stolle *et al*. 2019; Brelsford *et al*. 2020). But not all supergenes are necessarily expected to degenerate. In *SD-Mal*, for instance, complementing *SD-Mal* haplotypes can recombine via crossing over, if rarely, and gene flow from wildtype *SD*^+^ to *SD-Mal* chromosomes can occur via gene conversion. In the mouse *t*-haplotype, there is similar evidence for occasional recombination between complementing *t*-haplotypes (Dod *et al*. 2003) and with standard chromosomes, probably via gene conversion (Herrmann *et al*. 1987; Erhart *et al*. 2002; Wallace and Erhart 2008; Kelemen and Vicoso 2018). Despite the many parallels characterizing supergenes, their ultimate evolutionary fates depend on the particulars of the system.

## Material and methods

### Fly lines, library construction and sequencing

We sequenced haploid embryos using the scheme in Langely et al (Langley *et al*. 2011), which takes advantage of a mutation, *ms(3)K81* (Fuyama 1984), which causes the loss of the paternal genome early in embryonic development. We crossed *SD-Mal/CyO* stocks generated in (Brand *et al*. 2015) to homozygous *ms(3)K81* males and allowed them to lay eggs overnight. We inspected individual embryos under a dissecting scope for evidence of development and then isolated them for whole genome amplification using the REPLI-g Midi kit from Qiagen (catalog number 150043). For each WGA DNA sample, we tested for the presence of *Sd-RanGAP* using PCR (primers from (Presgraves *et al*. 2009). We prepared sequencing libraries for Illumina sequencing with TruSeq PCR free 350bp. We assessed library quality using a BioAnalyzer and sequenced with HiSeq2500 2×150bp reads (TruSeq) or 2×125bp reads (Nextera). To trim reads, we used Trimgalore v0.3.7 and the parameters: *q 28 --length 20 --paired -a GATCGGAAGAGCACACGTCTGAACTCCAGTCAC-a2 GATCGGAAGAGCGTCGTGTAGGGAAAGAGTGT --phred33 --fastqc --retain_unpaired -r1 21 -r2 21 --dont_gzip --length 20*. Trimmed reads are available in SRA (Bioproject PRJNA649752, SRA accession numbers in Table S1)

We sequenced a total of 10 *SD-Mal* genomes. One of these genomes (*SD-ZI157*) was *Sd-In(2R)Mal*^+^, non-driving, and therefore excluded from further analysis. Out of the remaining 9 driving *SD-Mal* chromosomes, one of them (*SD-ZI138*) had lower depth than the other 8 (Table S1, sheet 2) in the main chromosome arms but unusually high depth in the mitochondrial genome. We ran additional analyses dropping *SD-ZI138* and show that including this sample does not affect our main conclusions (Supp. Table S5; sheet 2).

For the Nanopore library, we extracted High-Molecular-Weight DNA from ~200 frozen female *SD-ZI125/SD-ZI125* virgins. We extracted DNA using a standard phenol-chloroform method and spooled DNA using capillary tubes. We constructed a library with ~1 ug DNA using RAD004 kit and the ultra-long read sequencing protocol (QUICK). We sequenced the library using R9.4 flow cells and called bases with the ONT Albacore Sequencing Pipeline Software version v2.2.10.

### Estimating Rsp copy number

We mapped Zambian *SD* reads to an assembly containing *2R* pericentric heterochromatin (Chang and Larracuente 2019), including the *Rsp* locus detailed in Khost et al. (Khost *et al*. 2017) with bowtie2 v2.3.5 (Langmead and Salzberg 2012). We estimated mean per-window and per-*Rsp* repeat depth using mosdepth v0.2.9 (Pedersen and Quinlan 2018). Coordinates for *Rsp* repeats were based on annotations in Khost et al. (2017).

### In(2R)Mal breakpoints

To assemble *SD-ZI125*, we filtered Nanopore reads using Porechop (v0.2.3) and Filtlong (--min_length 500) to remove adapters and short reads (https://github.com/rrwick/Porechop; (Wick *et al*. 2017) and https://github.com/rrwick/Filtlong). We were left with a total of 1,766,164,534 bases in 327,248 filtered reads. We performed *de novo* assemblies with the Nanopore reads using Flye v2.3.7 (Kolmogorov *et al*. 2019) with parameters “-t 24 -g 160m --nano-raw” and wtdbg v2.2 (Ruan and Li 2020) with parameters - p 19 -AS 1 -s 0.05 -L 0 -e 1”. We independently polished these two assemblies 10 times with Pilon v1.22 (Walker *et al*. 2014) using paired-end Illumina reads. Lastly, we reconciled these two polished assemblies using quickmerge v0.3 (Chakraborty *et al*. 2016) using the flye assembly as the reference with the command “python merge_wrapper.py wtdbg assembly flye assembly”. We aligned the contig containing the euchromatin on *SD-ZI125* to chromosome *2R* of the *D. melanogaster* (BDGP6) genome using Mauve (Darling *et al*. 2010). We defined the breakpoints according to the block rearrangement shown in Figure 1. To validate these breakpoints, we designed primers anchored at both sides of the most external breakpoints of *In(2R)Mal* (Suppl.Table S3) for PCR.

### Measuring genetic distances along SD-Mal and strength of distortion in the recombinants

To estimate recombination frequencies and obtain *SD* recombinant genotypes, we used a stock (*al[1] dpy[ov1] b[1] pr[1] c[1] px[1] speck[1]*, BDSC156, Bloomington Drosophila Stock Center), which has three visible, recessive markers on chromosome 2: *black (b*, 2L: 13.82), *curved* (*c*, 2R:15.9) and *plexus (px*, 2R:22.49). All crosses were transferred to a fresh vial after 5 days, and then adults were removed from the second vial after 5 days. Progeny emerging from the crosses were scored for up to 20 days following the cross.

To generate *SD-Mal* recombinant chromosomes, we crossed 8-10 *b,c,px/b,c,px* virgin females to 3-5 *SD-ZI125* males, recovered *SD-ZI125/b,c,px* virgins, then backcrossed 8-10 of them to 3-5 *b,c,px* homozygous males. To estimate genetic distance between the visible markers, we scored the number of recombinants in 11 crosses (n=1820). To compare genetic distance in *SD-Mal* to wild-type chromosomes, we estimated the number of recombinants from 15 crosses between *OregonR/b,c,px* females to *b,c,px/b,c,px* males (n=1716).

We recovered three types of recombinant chromosomes from *SD-ZI125/b,c,px* × *b,c,px/b,c,px* crosses: *b,Sd^+^,c^+^,In(2R)Mal,px*^+^; *b^+^,Sd,c,In(2R)Mal^+^,px* and *b^+^,Sd,c^+^,In(2R)Mal,px*. We crossed 3-5 virgin *b,c,px/b,c,px* females to individual recombinant males of each type, and scored the proportion of progeny carrying the recombinant chromosome (*k*=n+_recombinant_/n_total_). To distinguish distortion from viability effects, we also measured transmission of recombinant chromosomes through females, as drive is male-specific. We used these crosses to measure relative viability (*w*=n_recombinant_/n_*bcpx*_). We then used *w* to calculate a viability-corrected strength of distortion in males (*k\**=n_recombinant_/(*w*n_bcpx_+n_recombinant_) (Powers and Ganetzky 1991).

### Estimate of the frequency of SD-Mal in the DPGP3 dataset

To estimate the frequency of *In(2R)Mal* in a random sample of Zambian chromosomes, we mapped the 204 Illumina paired-end libraries from the DPGP3 dataset (Lack *et al*. 2016) to the *D. melanogaster* (BDGP6) genome, using bwa-mem (v0.7.9a), and we visually looked for an accumulation of discordant read pairs surrounding the estimated breakpoints of *In(2R)Mal*. To test the reliability of this method, we also applied it to detect cosmopolitan inversions *In(2L)t* and *In(2R)NS* and compared our inversion calls with the most recent inversion calls for the DPGP3 dataset (http://johnpool.net/Updated_Inversions.xls, last accessed 07/13/2020), getting a 98% and 99% of concordance for *In(2L)t* and *In(2R)NS*, respectively. To determine the frequency of the *Sd-RanGAP* duplication in the DPGP3 dataset we applied a similar method around the breakpoints of the *Sd-RanGAP* duplication (see Suppl.Table S4).

### SNP calling and annotation

For SNP calling, we mapped the Illumina reads from our *SD-Mal* libraries and the 20 *SD*^+^ libraries from the *DPGP3* dataset to *D. melanogaster* (BDGP6) genome (ftp://ftp.ensembl.org/pub/release-88/fasta/drosophila_melanogaster/dna/; last accessed 6/25/20) using BWA mem (v0.7.9a). We removed duplicated reads with Picard (2.0.1) and applied the GATK (3.5) “best practices” pipeline for SNP calling. We did local realignment and base score recalibration using SNPs data from DPGP1 ensembl release 88 (ftp://ftp.ensembl.org/pub/release-88/variation/vcf/drosophila_melanogaster/; last accessed 6/25/20). To call SNPs and indels, we used HaplotypeCaller and performed joint genotyping for each of the five genotypes using GenotypeGVCFs. SNPs filtered with following parameters: ‘QD < 2.0 || FS > 60.0 || MQ < 40.0 || MQRankSum < −12.5 || ReadPosRankSum < −8.0’. We annotated SNPs as synonymous or nonsynonymous using SNPeff (4.3, Cingolani *et al*. 2012) with the integrated *D. melanogaster* database (dmel_r6.12) database and parsed these annotations with SNPsift (Cingolani *et al*. 2012). To classify the SNPs as ‘shared’ between *SD-Mal, SD^+^In(2L)t* and *SD^+^In(2R)NS*, or ‘private’ to each one of them, we used BCFtools intersect (1.6; Danecek *et al*. 2021).

### Population genomics analysis

We wrote a Perl script to estimate *S*, *π*, Tajima’s *D*, *F*_ST_ and *d_XY_* in windows across the genome (available here: https://github.com/bnavarrodominguez/sd_popgen). To calculate *F*_ST_ values, we used the Weir-Cockerham estimator (Weir and Cockerham 1984). Only those sites with a minimum sample depth of 8 were included in the *F*_ST_ and Tajima’s *D* calculations. We determined window size by the number of ‘acceptable sample depth’ sites (and not, for example, a particular range of chromosome coordinates). To confirm that repeats were not interfering with our results, we ran our population genomics pipeline masking SNPs in repetitive elements identified by RepeatMasker (Smit *et al*. 2013), which yielded equivalent results (Suppl. Table S5, sheet 1).

### Age of the sweep

We calculated overall *S_In(2R)Mal,_ π_In(2R)Mal_* and Tajima’s *D_In(2R)Mal_* from the *SD-Mal* SNP set using our same Perl script (available here: https://github.com/bnavarrodominguez/sd_popgen), using a single window of 9.5Mb within the boundaries of *In(2R)Mal*. To account for gene conversion, we calculated an additional set of summary statistics masking the SNPs annotated as shared by at least one of the *SD*^+^ libraries. We estimated the time since the most recent selective sweep using an ABC method with rejection sampling. We modeled the selective sweep as an absolute bottleneck (*N_t_=1*) at some time (*t, 4N_e_* generations) in the past. We performed simulations in *ms* (Hudson 2002), considering a sample size of 9 and assuming no recombination in the ~9.92 Mb of *In(2R)Mal*. We simulated with values of *S_Sim_* drawn from a uniform distribution ±5% of *S_In(2R)Mal_*. We considered a prior uniform distribution of time of the sweep (*t*) ranging from 0 to *4N_e_* generations, *i.e*., 0 – 185,836 years ago, considering *D. melanogaster Ne* in Zambia 3,160,475 (Kapopoulou *et al*. 2018), frequency of *In(2R)Mal* 1.47% and 10 generations per year. The rejection sampling algorithm is as follows: (1) draw *S_Sim_* and *t* from prior distributions; (2) simulate 1000 samples using the coalescent under a selective sweep model; (3) calculate average summary statistics for drawn *S_Sim_* and t; (4) accept or reject chosen parameter values conditional on *|π_In(2R)Mal_* – *π_Sim_|* ≤ *ε*,|*D_In(2R)Mal_* – *D_Sim_*| ≤ *ε;* (5) return to step 1 and continue simulations until *m* desired samples from the joint posterior probability distribution are collected. For estimates of *t*, *ε* was set to 5% of the observed values of the summary statistics (in step 4) and *m* was set to 10,000. These simulations were performed with parameters calculated using all the SNPs in *In(2R)Mal* and excluding SNPs shared with *SD*^+^ chromosomes to account for gene conversion. We simulated 100,000 samples with the resulting estimated *t* and *S_In(2R)Mal_*, under our sweep model and under a constant size population model, and calculated two-sided *p* values for π and Tajima’s *D* using an empirical cumulative probability function (ecdf) in R (Team 2019). We estimated the maximum *a posteriori* estimate (MAP) as the posterior mode and 95% credibility intervals (CI) in R (Team 2019).

### Recombination

For estimates of recombination, we filtered the SNPs in *In(2R)Mal* to variable positions genotyped in all of the 9 *ZI-SD* samples and excluded singletons, resulting in a total of 338 SNPs. We estimated pairwise linkage disequilibrium (*r^2^*) using PLINK v1.9 (Purcell *et al*. 2007). We discarded *r^2^* data calculated for pairs of SNPs flanking the internal *In(2R)Mal* breakpoints. For comparison, we estimated pairwise linkage disequilibrium in the same region of *In(2R)Mal* for *SD*^+^ uninverted *2R* chromosome arms and, for comparison, in *SD^+^ In(2R)NS* inversion and in *SD^+^ In(2L)t* inversions. For *SD*^+^ chromosomes we applied the same filters (variable, non-singleton SNPs), plus a SNP ‘thinning’ to 1 SNPs/kb to get a manageable set of results. To investigate the possibility of crossing over between *SD-Mal* chromosomes, we used RecMin (Myers and Griffiths 2003) to estimate the minimum number of crossovers between the 338 bi-allelic, non-singleton SNPs in *In(2R)Mal*. RecMin input is a binary file, which we generated using *SD-ZI125* as an arbitrary reference for *SD*, assigning 0 or 1 on each position depending on if it was the same base or different. Maximum likelihood trees to establish relationships between *SD-Mal* haplotypes based on these 338 SNPs were estimated using RAxML-NG (Kozlov *et al*. 2019).

Runs of shared and private SNPs were identified in R, using all SNPs (including singletons). A run of SNPs is defined as a region from 5’ to 3’ where all the SNPs are in the same category (shared or private). Distance between the first and the last SNP of a category is considered length of the run. The region between the last SNP of a category and the first SNP of the alternative category is considered distance between runs. Because our sample size is small, we may underestimate the number of shared SNPs, as some private SNPs may be shared with some *SD*^+^ chromosomes that we have not sampled.

### Transposable element calling

We used a transposable element (TE) library containing consensus sequences of *Drosophila* TE families (Chang and Larracuente 2019). With this library, we ran RepeatMasker (Smit *et al*. 2013) to annotate reference TEs in the *D. melanogaster* (BDGP6) genome. To detect genotype specific TE insertions in our Illumina libraries, we used the McClintock pipeline (Nelson *et al*. 2017), which runs six different programs with different strategies for TE calling. We collected the redundant outputs from RetroSeq (Keane *et al*. 2013), PoPoolationTE (Kofler *et al*. 2012), ngs_te_mapper (Linheiro and Bergman 2012), TE-Locate (Platzer *et al*. 2012) and TEMP (Zhuang *et al*. 2014), discarded the calls produced by TEMP based on non-evidence of absence, and then merged the insertions detected by all different programs, considering the same insertion those of the same TE closer than a distance of +/- 600 bp, as described in (Bast *et al*. 2019). To reduce false positives, we only considered TE insertion calls that were predicted by more than one of the methods. To account for differences in library read number and/or length between datasets, we report the TE counts for *2R* normalized by the TE count for chromosome *2L* for the same library (Figure 6b). To assess whether library differences qualitatively affect our results, we repeated the above TE analysis on a set of 3 million randomly selected paired-end reads, trimmed to a fixed length of 75 bp, from each library and report TE count for chromosomes *2R* and *2L* separately (Suppl. Fig. S7).

## Acknowledgements

This work was funded by the National Institutes of Health (NIH) National Institute of General Medical Sciences (R35GM119515 and NIH-NRSA F32GM105317to A.M.L.), Stephen Biggar and Elisabeth Asaro fellowship in Data Science to A.M.L, a David and Lucile Packard Foundation grant and University of Rochester funds to D.C.P. We thank Dr. Danna Eickbush for assistance with genomic DNA preparation for nanopore sequencing. We also thank the University of Rochester CIRC for access to computing cluster resources and UR Genomics Research Center for the library construction and sequencing.

## Data availability

Raw sequence data are deposited in NCBI’s short read archive under project accession PRJNA649752. All code for data analysis and figure generation is available in Github (https://github.com/bnavarrodominguez/sd_popgen).

## Supplemental figure legends

Suppl. Fig. S1. Estimated abundance of *Rsp* repeats at each *Rsp* locus in the reference *Iso-1* genome and *SD-Mal*. For each locus annotated in the reference *D. melanogaster* genome (Khost *et al*. 2017), we plot estimated *Rsp* abundance as the sum of average depth of repeats at each locus normalized by average depth of chromosome 2 on the y-axis. *SD-Mal* has very few reads mapping to the primary *Rsp* locus (*Rsp-proximal* and *Rsp-major*), suggesting a complete deletion of the target of drive.

Suppl. Fig. S2. Model of the *In(2R)Mal* rearrangement. (a) Wild-type arrangement of chromosome *2R*. Pericentromeric heterochromatin and the centromere are represented by a grey rectangle and black circle, respectively. (b) *In(2R)Mal-d*: inversion of 4.18 Mb of 2R (*2R*:14,591,003-18,774,475), which disrupted the 3’ UTR of the *Mctp* gene (*2R*:18,761,758 - 18,774,824). (c) *In(2R)Mal-p*: Inversion of 6.76 Mb of 2R (*2R*:8,855,602-17,749,310), with 1.02 Mb overlapping with the now proximal segment of *In(2R)Mal-d*. This inversion disrupted the 3’ UTR of the *sns* gene (*2R*:8,798,489 - 8,856,091) and the CDS of the *CG10931* gene (*2R*:17,748,935 −17,750,136).

Suppl. Fig. S3. Average pairwise nucleotide diversity per site (π) in non-overlapping, 1-kb windows, for *SD*^+^ and *SD-Mal* chromosomes, around the *Sd-RanGAP* locus.

Suppl. Fig. S4. Neutral coalescent simulations under a constant size population model and a sweep and expansion model (absolute bottleneck at a time *t*, between 0 and 4*N_e_* generations). Simulations were done with *S* estimated using all SNPs in *In(2R)Mal* (a) and excluding SNPs shared with *SD*^+^ chromosomes (b) to account for gene conversion. Blue horizontal line marks observed *π_In2RMal_* (estimated for the entire region) and Tajima’s *D* in *In(2R)Mal* in Zambian *SD* chromosomes.

Suppl. Fig. S5. Frequency spectra of synonymous and non-synonymous SNPs in the *In(2R)Mal* chromosome region, in Zambian *SD* chromosomes (n=9, orange) and wildtype chromosomes from the same population, bearing the cosmopolitan inversions *In(2L)t* (n=10, dark blue) and *In(2R)NS* (n=10, light blue). *N/S* ratio for each of the frequency categories.

Suppl. Fig. S6. Number of insertions per TE family in *SD-Mal* compared to uninverted SD^+^ chromosome *2R*, both in *In(2R)Mal (2R:8.85-18.77*, top panel) and the region distal to it (*2R:18.77-25.29*, bottom panel). The families *DNA/M4DM, LTR/MDG1, LTR/ROO_I* and *Non-LTR/LINE* are highly overrepresented in *In(2R)Mal*.

Suppl. Fig. S7 Abundance of TEs in down sampled (3M reads, 75-bp) libraries for Zambian *SD* chromosomes (n=9, orange) and *SD*^+^ chromosomes from the same population, bearing the cosmopolitan inversions *In(2L)t* (N=10, dark blue) and *In(2R)NS* (N=10, light blue).

